# Antigenicity assessment of SARS-CoV-2 saltation variant BA.2.87.1

**DOI:** 10.1101/2024.03.07.583823

**Authors:** Sijie Yang, Yuanling Yu, Fanchong Jian, Ayijiang Yisimayi, Weiliang Song, Jingyi Liu, Peng Wang, Yanli Xu, Jing Wang, Xiao Niu, Lingling Yu, Yao Wang, Fei Shao, Ronghua Jin, Youchun Wang, Yunlong Cao

## Abstract

The recent emergence of a SARS-CoV-2 saltation variant, BA.2.87.1, which features 65 spike mutations relative to BA.2, has attracted worldwide attention. In this study, we elucidate the antigenic characteristics and immune evasion capability of BA.2.87.1. Our findings reveal that BA.2.87.1 is more susceptible to XBB-induced humoral immunity compared to JN.1. Notably, BA.2.87.1 lacks critical escaping mutations in the receptor binding domain (RBD) thus allowing various classes of neutralizing antibodies (NAbs) that were escaped by XBB or BA.2.86 subvariants to neutralize BA.2.87.1, although the deletions in the N-terminal domain (NTD), specifically 15-23del and 136-146del, compensate for the resistance to humoral immunity. Interestingly, several neutralizing antibody drugs have been found to restore their efficacy against BA.2.87.1, including SA58, REGN-10933 and COV2-2196. Hence, our results suggest that BA.2.87.1 may not become widespread until it acquires multiple RBD mutations to achieve sufficient immune evasion comparable to that of JN.1.

---

Recently, BA.2.87.1 gained wide attention for its expansion with striking number of spike mutations^1,2^. This novel saltation variant harbors 65 mutations on the Spike glycoprotein, notably including two unique extensive NTD deletions 15-26del and 136-146del, and RBD substitutions such as K417T, K444N, V445G, L452M, and N481K (Figure 1A). With eight sequences originating from South Africa in the last quarter of 2023 and one travel-related sequence in USA, it was officially designated in February 2024 and has then been tracked by USA CDC (Centers for Disease Control and Prevention)^3^. Some of the prior saltation variants of SARS-CoV-2 have demonstrated profound capability of evading humoral immunity induced by vaccinations and infections due to the substantial mutations and antigenic difference, with the most notable examples Omicron BA.1, BA.2.75 and the recent JN.1^4–17^ (Figure 1B). Consequently, it is imperative to rapidly assess BA.2.87.1 potential increases in immune resistance.

**Figure 1.**
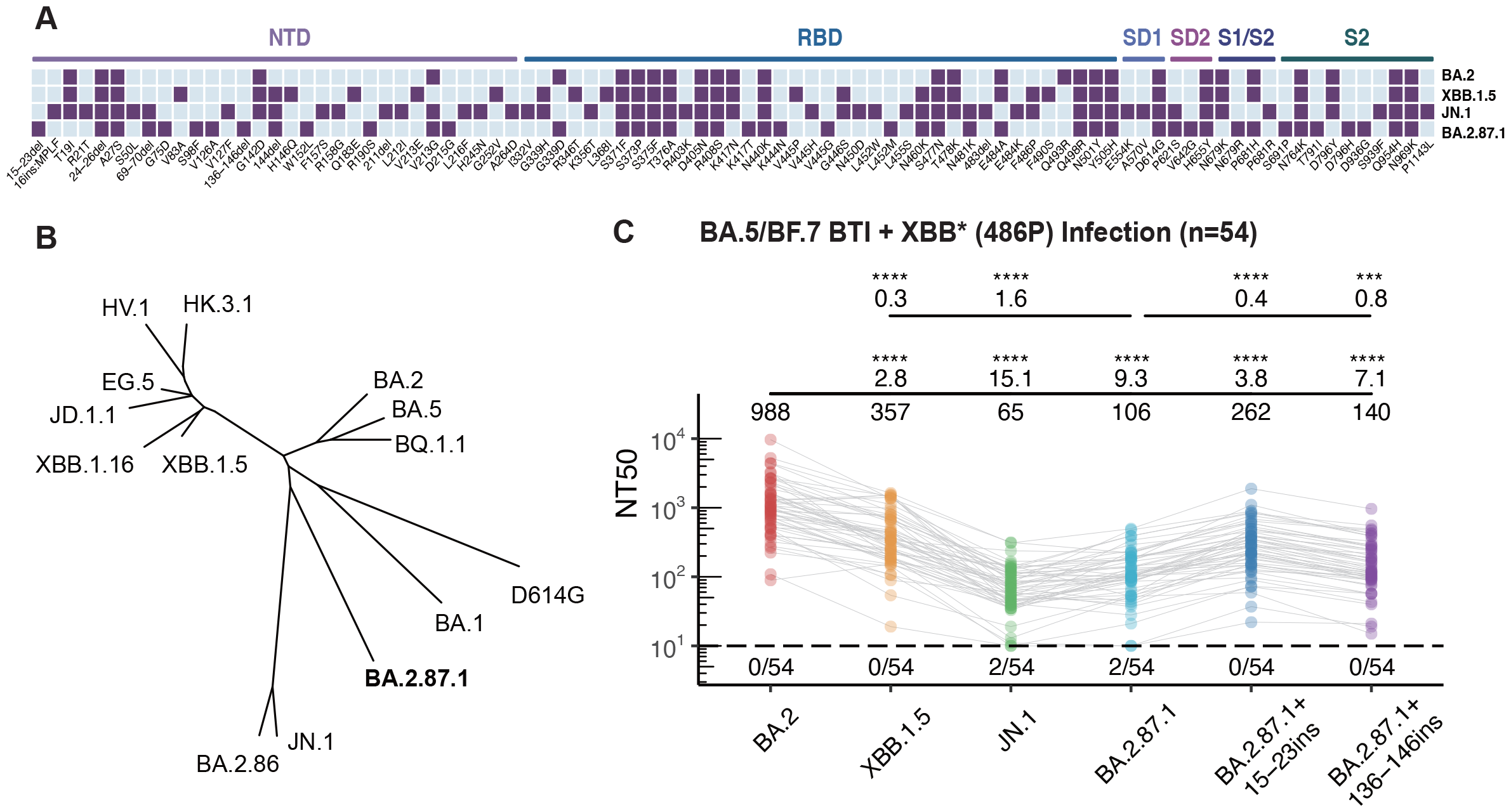
Spike mutations, phylogeny and plasma neutralization resistance of BA.2.87.1. (A) Mutations on the spike glycoprotein of BA.2, XBB.1.5, JN.1, and BA.2.87.1. Purple square indicates the presence of the mutations in each variant, while sky blue square indicates the absence of the mutations. The domains of the mutations on the Spike protein are annotated above. (B) Unrooted phylogenetic tree of the Spike glycoprotein of prevalent SARS-CoV-2 variants. (C) The 50% neutralizing titers (NT50) of convalescent plasma from individuals who experienced breakthrough infections with BA.5 or BF.7 followed by reinfection with XBB*+486P subvariants (n=54) against SARS-CoV-2 variants. Geometric mean titers, relative fold changes, and statistical significance are annotated above each group. The dashed line represents the limit of detection (50% neutralizing titer=10), and the number of negative samples is indicated below it. Paired samples were analyzed by two-tailed Wilcoxon signed-rank test. ***p< 0.001, ****p< 0.0001.

First, to investigate BA.2.87.1’s potential for humoral immune escape, vesicular stomatitis virus (VSV)-based pseudovirus neutralization assays were performed on plasma samples obtained from participants (n=54) who underwent BA.5/BF.7 breakthrough infection (BTI) after three doses of ancestral-strain inactivated vaccination, and then re-infected with XBB*+486P (Figure 1C and Table S1). The results suggested that pseudovirus bearing the BA.2.87.1 spike protein demonstrated superior evasion to XBB.1.5, indicated by a 70% percent decrease in 50% neutralizing titers (NT50). However, it exhibited lower ability to evade humoral immunity induced by BA.5/BF.7 BTI and XBB reinfection when compared to JN.1, with a 1.6-fold higher NT50. To evaluate the impact of the two long NTD deletions on plasma immune escape, two NTD segments of BA.2 (the precursor of BA.2.87.1) on the corresponding residues 15-23 (CVNLITRTQ) and 136-146 (CNDPFLDVYYH) were inserted into BA.2.87.1 pseudovirus and tested for plasma neutralization. Both the reversion of 15-23del and 136-146del significantly dampen the antibody evasion of BA.2.87.1, with 2.5-fold and 1.3-fold increase in NT50, respectively. In summary, neutralization experiments with human plasma revealed that NTD deletion profoundly contributed to the plasma escape of BA.2.87.1, allowing it to exhibit higher immune evasion capability than BA.2 and XBB.1.5. However, its overall immune evasion strength was not comparable to that of JN.1.

Subsequently, we evaluated the neutralizing potency of a collection of BA.2-effective monoclonal antibodies (mAbs) derived from repetitive Omicron infections that target various epitopes of RBD, as defined in our previous studies, to further interrogate BA.2.87.1’s immune escape capability and mechanism (Figure 2A)^4,18–21^. Most of the mAbs involved remain reactive to XBB.1.5 as well. We found that BA.2.87.1 could be easily neutralized by most of the antibodies in the A1, A2, B and F3 epitope groups (Class 1, Class 2 and Class 1/4 antibodies), which directly compete with the binding of host angiotensin-converting enzyme 2 (ACE2) to realize potent inhibition^7,21^. In contrast, the previously and recently circulating evasive variants, including HK.3.1, JD.1.1 and JN.1, could efficiently escape these antibodies due to their escape mutations on the Spike receptor-binding motif (RBM). Specifically, the “FLip” mutation (L455F + F456L) of HK.3.1 improved its ability to evade A1 and A2 antibodies, and the additional A475V mutation carried by JD.1.1 further enhances the evasion^22^. Furthermore, the lack of F486P but the achievement of N481K mutation of BA.2.87.1 compared to XBB.1.5 lineages demonstrated its relatively weak yet distinct escape pattern against Group B antibodies^23^. Without the R346T and K356T mutations, BA.2.87.1 could not efficiently evade Class 3 antibodies (Group D3, D4, E1/E2.1)^24^. In addition, JN.1+R346T showed slightly enhanced resistance to antibodies from Class 3. These observations suggest that BA.2.87.1, despite being antigenically distinct, typically demonstrates less resistance to various classes of RBD-targeting antibodies compared to JN.1.

**Figure 2.**
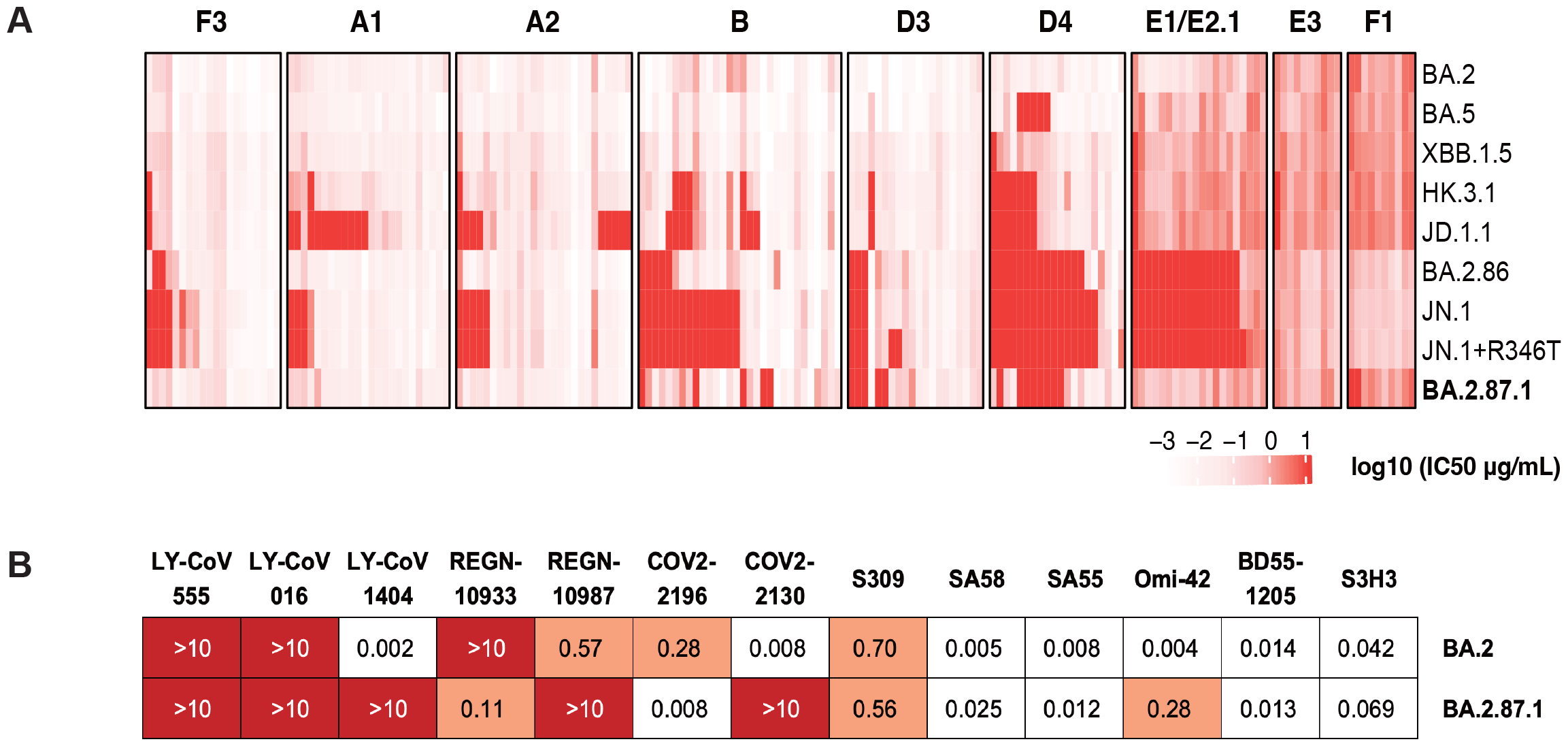
Neutralization of monoclonal antibodies against BA.2.87.1. (A) The 50% inhibitory concentration (IC50, μg/mL) of a panel of BA.2-effective monoclonal neutralizing antibodies targeting distinct RBD epitopes (determined by DMS) against BA.2, BA.5, XBB.1.5, HK.3.1, JD.1.1, BA.2.86, JN.1, JN.1+R346T, and BA.2.87.1 mutants. Epitope groups are annotated above. The intensity of red represents the magnitude of the IC50 values. (B) IC50 (μg/mL) of approved or candidate monoclonal neutralizing antibody drugs targeting the RBD or SD1 regions of the spike protein, against BA.2 and BA.2.87.1 pseudoviruses.

As for the therapeutic antibodies, SA55 maintained consistent effectiveness against BA.2.87.1 (Figure 2B)^25^. REGN-10933 and COV2-2196, which target the B epitope, regained their neutralizing capacity against BA.2.87.1, likely due to the absence of mutations on F486, which was usually mutated to Val, Ser, or Pro in existing BA.5, XBB.1.5, and BA.2.86 subvariants^26,27^. Similarly, the SA58 antibody also restored its potency, attributable to the lack of G339H and R346T mutations^25^. Moreover, BA.2.87.1 was also unable to resist the SD1-targeting S3H3 without the E554K mutation which was carried by the BA.2.86 lineages^28^. Nevertheless, due to the K444N and L452M mutations, BA.2.87.1 exhibited robust resistance to Group D1/D2 antibodies, such as REGN-10987, LY-CoV1404, and COV2-2130^26,29,30^. As for Class 1 broad-spectrum antibodies, Omi-42 exhibited substantially compromised neutralization, while BD55-1205, which belonged to the public IGHV3-53/3-66 NAb group, maintained its neutralization potency against all tested variants^25,31^.

## Discussion

Consistent with the recent research^1^, our findings highlight that the NTD deletions, particularly the 15-23del, significantly contribute to the evasion of BA.2.87.1. Such long stretches of NTD deletions are frequently observed in persistent evolution, underscoring the need to monitor the intra-host persistent evolution of SARS-CoV-2^32–34^. We also observed a partial recovery of therapeutic antibodies^2^. However, it is highlighted that BA.2.87.1 demonstrates weaker immune resistance to the currently dominant variant JN.1. Our results also emphasized that BA.2.87.1 cannot counteract Group A and B antibodies without mutations such as L455S, L455F, F456L, A475V and F486P, which are present in JN.1 and JD.1.1. Furthermore, in the absence of mutation at position R346 or K356, its exhibit weak capability of escape NAbs in the E1/E2.1 epitope group. These characteristics collectively suggest that BA.2.87.1 is not likely to outcompete current JN.1 and XBB subvariants without substantially further antigenic drift on the RBD, given the current population-level immunity established by the repeated vaccinations and infections. Although BA.2.87.1 is relatively weak in terms of antigenicity, its other virological characteristics and potential of accumulating additional RBD mutations during regional circulation should be closely evaluated and monitored.

## Acknowledgments

We extend our gratitude to scientists in the community for their persistent monitoring of SARS-CoV-2 variants and valuable discussion. We thank all volunteers who contributed blood samples for this research. This project is financially supported by the Ministry of Science and Technology of China (2023YFC3041500; 2023YFC3043200), Changping Laboratory (2021A0201; 2021D0102), and the National Natural Science Foundation of China (32222030, 2023011477).

## Author Contributions

Y.C. designed and supervised the study. S.Y. and Y.C. wrote the manuscript with inputs from all authors. Y.X. and R.J. recruited the SARS-CoV-2 convalescents. S.Y., F.J., J.L. and W.S. performed sequence analysis and illustration. Y.Y. and Youchun W. constructed pseudoviruses. P.W., L.Y., Yao W., J.W. and F.S. processed the plasma samples and performed the pseudovirus neutralization assays. S.Y., F.J., W.S., A.Y., X.N., and Y.C. analyzed the neutralization data.

## Declaration of interests

Y.C. is the inventor of the provisional patent applications for BD series antibodies, which includes BD55-5514 (SA55), BD55-5840 (SA58) and BD55-1205. Y.C. is the founder of Singlomics Biopharmaceuticals. Other authors declare no competing interests.

## Methods

### Plasma isolation

Blood samples were obtained from individuals who had recovered from or been re-infected with the SARS-CoV-2 Omicron BTI variant, following the research protocol approved by Beijing Ditan Hospital, affiliated with Capital Medical University (Ethics Committee archiving No. LL-2021-024-02), the Tianjin Municipal Health Commission, and the Ethics Committee of Tianjin First Central Hospital (Ethics Committee archiving No. 2022N045KY). Prior to sample collection, all participants provided written informed consent, consenting to the collection, storage, and utilization of their blood samples solely for research purposes, as well as the subsequent publication of associated data. Patients in the reinfection group were initially infected with the BA.5/BF.7 variants in December 2022 in Beijing and Tianjin, China^35^. From December 1, 2022, to February 1, 2023, over 98% of the sequenced samples obtained were identified as BA.5* (excluding BQ*), primarily consisting of the subtypes BA.5.2.48* and BF.7.14*, which typically represented the BA.5/BF.7 variants during this period. Subsequently, patients in the XBB BTI cohort and those with secondary infections in the reinfection group were infected between May and June 2023. Over 90% of the sequencing samples from Beijing and Tianjin during this period corresponded to the XBB*+486P variant. These infections were confirmed by polymerase chain reaction (PCR) or antigen testing.

Whole blood was diluted 1:1 with a solution of phosphate-buffered saline (PBS) supplemented with 2% fetal bovine serum (FBS). Following this, Ficoll gradient centrifugation (Cytiva, 17-1440-03) was performed. After centrifugation, the plasma was collected from the upper layer. The plasma samples were then stored in aliquots at 20°C or below and were heat-inactivated before subsequent experiments.

### Pseudovirus preparation

SARS-CoV-2 spike protein pseudovirus was produced based on vesicular stomatitis virus (VSV) pseudovirus packaging system^36^. To enhance the spike protein’s expression efficiency in mammalian cells, the spike protein gene was codon-optimized and inserted into the pcDNA3.1 expression plasmid, utilizing the BamHI and XbaI restriction enzyme sites. During pseudovirus production, the SARS-CoV-2 spike protein expression plasmid was transfected into 293T cells(American Type Culture Collection (ATCC, CRL-3216). Following transfection, the 293T cells were infected with G*ΔG-VSV virus (VSV-G pseudotyped virus, Kerafast) in the cell culture supernatant. The pseudovirus was then collected from the supernatant, harvested and filtered. The collected pseudovirus was aliquoted and stored at -80°C for future use.

### Pseudovirus neutralization assays

Pseudovirus neutralization assays were conducted utilizing the Huh-7 cell(Japan Collection of Research Bioresources [JCRB], 0403). Plasma samples or monoclonal antibodies were serially diluted and then mixed with pseudovirus. After incubation at 37°C with 5% CO2 for 1 hour, digested Huh-7 cells were added and incubated for 24 hours at 37°C. The supernatant was discarded, and the mixture was incubated with D-Luciferin reagent (PerkinElmer, 6066769) for 2 minutes in the dark. The cell lysate was then transferred to a detection plate, and luminescence intensity was measured using a microplate spectrophotometer (PerkinElmer, HH3400). IC50 values were determined using a four-parameter logistic regression model^37^.

## Supplementary Tables

**Table S1** | **Details of SARS-CoV-2 convalescent individuals involved in the study**

**Table S2** | **Sequences information of the constructed SARS-CoV-2 pseudoviruses**

## References

1. Wang, X. et al. Robust Neutralization of SARS-CoV-2 Variants Including JN.1 and BA.2.87.1 by Trivalent XBB Vaccine-Induced Antibodies. Preprint at doi:10.1101/2024.02.16.580615 (2024).

2. Zhang, L. et al. 1 Virological traits of the SARS-CoV-2 BA.2.87.1 lineage. Preprint at doi:10.1101/2024.02.27.582254.

3. CDC Tracking BA.2.87.1, New Omicron Subvariant With Potential to Evade Immunity | Vaccination | JAMA | JAMA Network. https://jamanetwork.com/journals/jama/article-abstract/2815864.

4. Cao, Y. et al. Omicron escapes the majority of existing SARS-CoV-2 neutralizing antibodies. Nature 602, 657–663 (2022).

5. Cao, Y. et al. Characterization of the enhanced infectivity and antibody evasion of Omicron BA.2.75. Cell Host Microbe 30, 1527-1539.e5 (2022).

6. Yang, S. et al. Antigenicity and infectivity characterisation of SARS-CoV-2 BA.2.86. Lancet Infect. Dis. 23, e457–e459 (2023).

7. Yang, S. et al. Fast evolution of SARS-CoV-2 BA.2·86 to JN.1 under heavy immune pressure. Lancet Infect. Dis., (2023).

8. Wang, Q. et al. Antigenicity and receptor affinity of SARS-CoV-2 BA.2.86 spike. Nature 1–3 (2023) doi:10.1038/s41586-023-06750-w.

9. Wannigama, D. L. et al. Tracing the new SARS-CoV-2 variant BA.2.86 in the community through wastewater surveillance in Bangkok, Thailand. Lancet Infect. Dis. 23, e464–e466 (2023).

10. Wang, Q. et al. Evolving antibody evasion and receptor affinity of the Omicron BA.2.75 sublineage of SARS-CoV-2. iScience 26, 108254 (2023).

11. Markov, P. V. et al. The evolution of SARS-CoV-2. Nat. Rev. Microbiol. 21, 361–379 (2023).

12. Addetia, A. et al. Neutralization, effector function and immune imprinting of Omicron variants. Nature 621, 592–601 (2023).

13. Sheward, D. J. et al. Sensitivity of the SARS-CoV-2 BA.2.86 variant to prevailing neutralising antibody responses. Lancet Infect. Dis. 23, e462–e463 (2023).

14. Uriu, K. et al. Transmissibility, infectivity, and immune evasion of the SARS-CoV-2 BA.2.86 variant. Lancet Infect. Dis. 23, e460–e461 (2023).

15. Sheward, D. J. et al. Evasion of neutralising antibodies by omicron sublineage BA.2.75. Lancet Infect. Dis. 22, 1421–1422 (2022).

16. Qu, P. et al. Immune evasion, infectivity, and fusogenicity of SARS-CoV-2 BA.2.86 and FLip variants. Cell, (2024).

17. Kaku, Y. et al. Virological characteristics of the SARS-CoV-2 JN.1 variant. Lancet Infect. Dis., (2024).

18. Yisimayi, A. et al. Repeated Omicron exposures override ancestral SARS-CoV-2 immune imprinting. Nature 625, 148–156 (2024).

19. Dejnirattisai, W. et al. The antigenic anatomy of SARS-CoV-2 receptor binding domain. Cell 184, 2183-2200.e22 (2021).

20. Piccoli, L. et al. Mapping Neutralizing and Immunodominant Sites on the SARS-CoV-2 Spike Receptor-Binding Domain by Structure-Guided High-Resolution Serology. Cell 183, 1024-1042.e21 (2020).

21. Barnes, C. O. et al. SARS-CoV-2 neutralizing antibody structures inform therapeutic strategies. Nature 588, 682–687 (2020).

22. Jian, F. et al. Convergent evolution of SARS-CoV-2 XBB lineages on receptor-binding domain 455–456 synergistically enhances antibody evasion and ACE2 binding. PLOS Pathog. 19, e1011868 (2023).

23. Cao, Y. et al. BA.2.12.1, BA.4 and BA.5 escape antibodies elicited by Omicron infection. Nature 608, 593–602 (2022).

24. Jian, F. et al. Further humoral immunity evasion of emerging SARS-CoV-2 BA.4 and BA.5 subvariants. Lancet Infect. Dis. 22, 1535–1537 (2022).

25. Cao, Y. et al. Rational identification of potent and broad sarbecovirus-neutralizing antibody cocktails from SARS convalescents. Cell Rep. 41, 111845 (2022).

26. Copin, R. et al. The monoclonal antibody combination REGEN-COV protects against SARS-CoV-2 mutational escape in preclinical and human studies. Cell 184, 3949-3961.e11 (2021).

27. Zost, S. J. et al. Potently neutralizing and protective human antibodies against SARS-CoV-2. Nature 584, 443–449 (2020).

28. Xu, S. et al. Mapping cross-variant neutralizing sites on the SARS-CoV-2 spike protein. Emerg. Microbes Infect. 11, 351–367 (2022).

29. Westendorf, K. et al. LY-CoV1404 (bebtelovimab) potently neutralizes SARS-CoV-2 variants. Cell Rep. 39, 110812 (2022).

30. Zost, S. J. et al. Potently neutralizing and protective human antibodies against SARS-CoV-2. Nature 584, 443–449 (2020).

31. Dijokaite-Guraliuc, A. et al. Rapid escape of new SARS-CoV-2 Omicron variants from BA.2-directed antibody responses. Cell Rep. 42, 112271 (2023).

32. McCallum, M. et al. N-terminal domain antigenic mapping reveals a site of vulnerability for SARS-CoV-2. Cell 184, 2332-2347.e16 (2021).

33. Díaz, Y. et al. SARS-CoV-2 reinfection with a virus harboring mutation in the Spike and the Nucleocapsid proteins in Panama. Int. J. Infect. Dis. 108, 588–591 (2021).

34. Weigang, S. et al. Within-host evolution of SARS-CoV-2 in an immunosuppressed COVID-19 patient as a source of immune escape variants. Nat. Commun. 12, 6405 (2021).

## Methods References

Pan, Y. et al. Characterisation of SARS-CoV-2 variants in Beijing during 2022: an epidemiological and phylogenetic analysis. The Lancet 401, 664–672 (2023).

Li, Q. et al. The Impact of Mutations in SARS-CoV-2 Spike on Viral Infectivity and Antigenicity. Cell 182, 1284-1294.e9 (2020).

Nie, J. et al. Establishment and validation of a pseudovirus neutralization assay for SARS-CoV-2. Emerg. Microbes Infect. 9, 680–686 (2020).

